# Multiplexed inhibition of immunosuppressive genes with Cas13d for on-demand combinatorial cancer immunotherapy

**DOI:** 10.1101/2023.03.14.532668

**Authors:** Feifei Zhang, Wang Guangchuan, Ryan Chow, Emily He, Medha Majety, Yueqi Zhang, Sidi Chen

## Abstract

Checkpoint blockade immunotherapy is a potent class of cancer treatment, however, the complex immunosuppressive tumor microenvironment (TME) often requires multi-agent combinations to be effective. Current cancer immunotherapy combination approaches are cumbersome, usually involving one-drug-at-a-time scheme. Here, we devise Multiplex Universal Combinatorial Immunotherapy via Gene-silencing (MUCIG), as a versatile approach for combinatorial cancer immunotherapy. We harness CRISPR-Cas13d to efficiently target multiple endogenous immunosuppressive genes on demand, allowing us to silence various combinations of multiple immunosuppressive factors in the TME. Intratumoral AAV-mediated administration of MUCIG (AAV-MUCIG) elicits significant anti-tumor activity with several Cas13d gRNA compositions. TME target expression analysis driven optimization led to a simplified off-the-shelf MUCIG targeting a four gene combination (PGGC: *Pdl1, Galectin9, Galectin3* and *Cd47*). AAV-PGGC shows significant *in vivo* efficacy in syngeneic tumor models. Single cell and flow profiling revealed that AAV-PGGC remodeled the TME by increasing CD8^+^ T cell infiltration and reducing myeloid-derived immunosuppressive cells (MDSCs). MUCIG thus serves as a universal method to silence multiple immune genes *in vivo,* and can be delivered via AAV as a therapeutic approach.

## Main

Cancer cells engage a variety of pathways to mold an immunosuppressive tumor microenvironment (TME) that favors tumor progression and therapy resistance ^1–3^. This immunosuppressive TME is often initiated from the primary tumor, subsequently evolving into a network of interlocking immunosuppressive mechanisms ^1, 4, 5^. For instance, tumors hyperactivate immune checkpoints to attenuate the effectiveness of T cells, allowing tumors to escape immune surveillance, suppress anti-tumor immunity, and hamper effective anti-tumor immune responses ^6^. By targeting these key inhibitory receptors, immune checkpoint blockade (ICB) therapy can unleash anti-tumor T cell responses ^7, 8^. In particular, PD-1/PD-L1 and CTLA-4 blockade have demonstrated significant clinical benefit in multiple tumor types ^9^. However, many patients do not respond to single agent or even combination ICB therapy. Consistent with their distinct mechanisms of action, concurrent combination therapy with anti-PD-1/PD-L1 plus anti-CTLA-4 appears to increase response rates above that of the corresponding monotherapies ^10–12^. Novel immunotherapy combinations may further increase the proportion of patients who respond to ICB. One major factor that limits the efficacy of ICB is that the immunosuppressive TME is highly dynamic and unique to each patient, even for patients with the same cancer type ^13^. Given the complexity and heterogeneity of the TME, targeting a single gene alone is often insufficient to provide clinical benefit to a broad range of patients. However, current immune checkpoint therapies primarily rely on monoclonal antibodies, blocking one target at a time using one antibody molecule each. Two or more antibodies have been used in combination; however, the difficulties for the approach of combining more and more antibodies scale exponentially, as development of each specific and potent therapeutic antibody is a daunting task by itself. A more flexible, versatile, and effective method for combinatorial immunotherapy is urgently needed.

Gene silencing offers a universal approach for reducing the expression of virtually any genes in the mammalian genome. Gene silencing methods include RNA interference (RNAi), CRISPR interference (CRISPRi), and more recently other RNA-targeted CRISPR effectors ^14, 15^. Simultaneous silencing of multiple genes has been readily achievable by multiplexing of target-specific guide sequences, such as short-hairpin RNAs (shRNAs), or CRISPR guide RNAs (gRNAs), as all genes’ silencing use the same mechanism and the same backbone machinery with these methods. Therefore, we reasoned that gene silencing may provide unique benefit for a substantially simpler, and more versatile approach for multiplexing targeting of immune genes. With the versatility of gene silencing, we further hypothesized that targeting multiple immunosuppressive genes in the TME in combinations would elicit anti-tumor immunity.

Here, we developed <u>M</u>ultiplex <u>U</u>niversal <u>C</u>ombinatorial <u>I</u>mmunotherapy via <u>G</u>ene-silencing (MUCIG), as a versatile method for combinatorial cancer immunotherapy. The recently discovered CRISPR/Cas13 systems have been demonstrated as efficient tools for RNA knockdown, as Cas13 proteins can bind and cleave endogenous RNAs in a programmable manner through the use of sequence-specific gRNAs ^15^. RfxCas13d (also known as CasRx) was reported as one such RNA targeting tool ^16, 17^, with a compact size of 990 amino acids that is considerably smaller than Cas13a-c effectors or Cas9, making it feasible to package both gRNAs and Cas13d into a single Adeno-associated virus (AAV) construct ^16^. Moreover, Cas13d is immunogenic, the antigen of which could induce T cell proliferative responses and increase CD4+T cell secreting IFN-γ or TNF-α ^18^, which could be utilized to enhance the tumor immune response. Harnessing CRISPR/Cas13d-based RNA targeting system for MUCIG, we set out to test the concept of combinatorial immunotherapy by simultaneously inhibiting multiple immunosuppressive genes in the TME *in vivo*.

### Efficient knockdown of endogenous immune suppressive genes using Cas13d

To assess the efficiency of Cas13d-mediated RNA knockdown of the endogenous immunosuppressive genes, 4 to 5 gRNAs for interested immunosuppressive genes: *Cd47, Galectin3* (*Lgals3*), *Cd66a* were designed based on the computational Cas13d gRNAs prediction tool^19^. We generated an all-in-one vector including gRNA, Cas13d and performed flow cytometry analyze gRNA knockdown efficacy by fluorescent intensity (**Figure S1A**). We found that the designed gRNAs could efficiently knock down the target genes (**Figure S1B**). For all 4 targeted genes, we identified at least one gRNA for each target gene that achieved over 50% knockdown efficiency. For *Cd66a*, all 5 designed gRNAs showed robust knockdown. These data indicated that the Cas13d gRNA design is predictive and reliable for the following multiplex genes targeting. We also compared the knockdown efficacy between Cas13d-mediated gRNAs and shRNAs, illustrating that Cas13d-mediated gRNA had better or similar knockdown than shRNAs, for the same genes in the same cell types (**Figure S1C&D).**To achieve stronger gene repression, we compared the knockdown efficiency of gRNAs bearing the wildtype direct repeat (WT-DR) vs. a mutant DR (Mut-DR) (**Figure 1A**), which was previously described to have improved efficiency ^19^. Flow cytometry analysis showed that the knockdown of PDL1 is less efficient in E0771 cells with the WT-DR g14 (∼11%). In contrast, the Mut-DR g14 could achieve ∼68% PDL1 knockdown at protein level (**Figure 1A**). For *Cd73* gRNA, we similarly observed improvement with Mut-DR. These data indicate that this modified Cas13d-gRNA system is an effective approach to repress the expression of tumor intrinsic immune suppressive genes.

**Figure 1.**
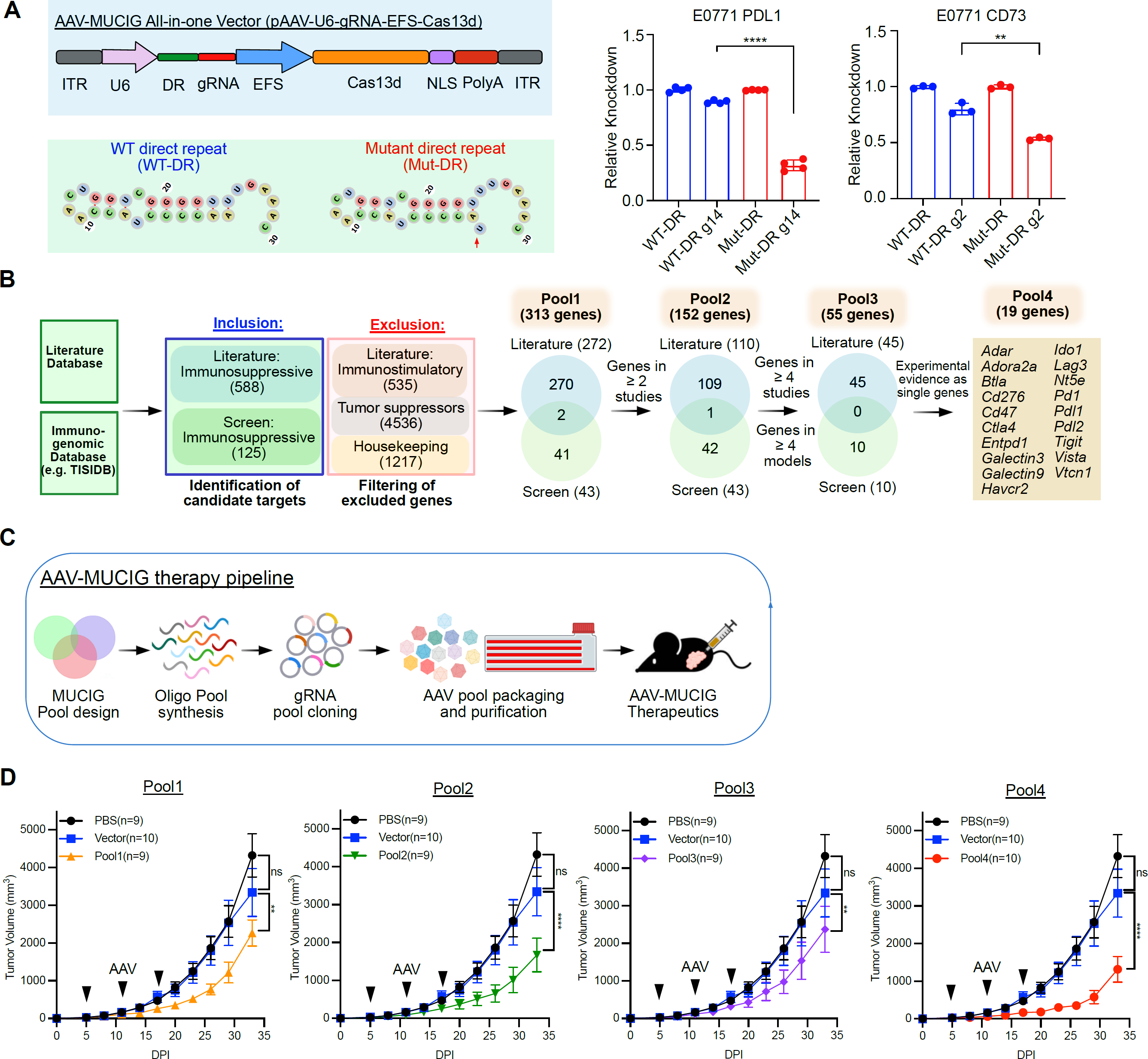
Multiplexed Cas13d repression of immunosuppressive genes as combinatorial cancer immunotherapy. **A.** Comparison of WT-DR (direct repeats) and mut-DR knockdown efficiency when targeting PDL1 and CD73 in E0771 cells. Data are expressed as the relative mean of fluorescent intensity (MFI). The gene expression level of WT-DR group was normalized to 1. Left top, design of an all-in-one AAV vector that contains an EFS-driven Cas13d expression cassette and a U6-driven Cas13d guide RNA cassette for repression test. Left bottom, WT-DR vs Mut DR structure. **B.** Design of four different gRNA libraries targeting immunosuppressive gene combinations. **C.** Schematics of the experimental design for evaluating <u>M</u>ultiplex <u>U</u>niversal <u>C</u>ombinatorial Immunotherapy via <u>G</u>ene-silencing (MUCIG) as immunotherapy. **D.** Growth curves of E0771 tumors in C57BL/6 mice. 2×10e^6^ E0771 cells were orthotopically injected into C57BL/6 mice. Mice were intratumorally injected with PBS (n = 9), AAV-MUCIG-Vector (Cas13d) (n = 10), AAV-MUCIG Pool1 (n = 9), Pool2 (n = 9), Pool3 (n = 9), or Pool4 (n = 10) at days 5, 9 and 14 with 2e^11^ AAV per dose. Data points in this figure are presented as mean ± s.e.m. Statistical significance was assessed by two-way ANOVA. ** *p* < 0.01, **** p < 0.0001. Non-significant comparisons not shown.

Recently, Cas13d was reported to have collateral activity in human cells ^20–22^. It was reported that when targeting the transfected DsRed in HEK cells, the co-transfected reporter gene GFP would be markedly down-regulated ^20^. However, when targeting the endogenous RNAs, the extent of collateral activity could be influenced by the abundance of the target RNA. To test how strong the collateral activity when targeting the endogenous immunosuppressive genes, we generated a GFP and mCherry dual reporter system to indicate the collateral activity of Cas13d (**Figure S2A**). Instead of transient transfection of the reporter gene plasmids, we established an E0771 cell line stably expressing Cas13d, GFP and mCherry protein by lentivirus transduction to better mimic the endogenous gene expression. According to the flow cytometry results, both the GFP and mCherry reporters showed stable expression among all the tested guide RNAs targeting the immunosuppressive genes, including non-transduced control (NTC) and empty vector (EV) (**Figure S2B**). Furthermore, we tested the specific gene targeting of these guide RNAs at protein by flow cytometry (**Figure S2B**). Even though scramble control caused a very mild background knockdown of PDL1 or GALECTIN9, the on-target knockdown is still much stronger (**Figure S2B**). Then RT-qPCR was performed to test the gene knockdown at RNA level. The data and statistical test among groups showed specific targeting of all the guide RNAs (**Figure S2C**). These data suggested specific on-target of the Cas13d guide RNAs when targeting the endogenous immunosuppressive genes.

### AAV-mediated immunosuppressive gene repression as an immunotherapeutic modality

Given that gene knockdown is not complete by Cas13d, the natural question is whether such degree of knockdown can lead to effective immune modulation, and thereby anti-tumor immunity *in vivo*. AAV is one of the leading vehicles for transgene delivery ^23^. To evaluate the feasibility of *in vivo* Cas13d and gRNA intratumoral delivery, we first generated an AAV vector expressing firefly luciferase and GFP (AAV-Luci-GFP). We intratumorally injected AAV-Luci-GFP into E0771 tumor-bearing mice and analyzed luciferase activity by *in vivo* bioluminescent imaging (**Figure S3A**). The time course imaging showed that luciferase was persistently expressed primarily in the tumor and, unsurprisingly, also in the liver (**Figure S3A).** These data indicate that intratumoral AAV injection can successfully deliver genetic cargo into tumor. Then, to evaluate which cell type(s) the intratumoral injection of AAV-Cas13d transduces, we administrated E0771-mCherry tumor with AAV-Luci-GFP or AAV-Cas13d-GFP (**Figure S3B**). Flow cytometry data analysis of the TME was performed by analyzing the GFP+ cells (**Figure S3C**). The data showed intratumoral injected AAV transduced both tumor cell and CD45+ immune cells (**Figure S3D**).

Having evaluated the feasibility of the Cas13d gRNA knockdown system, we next sought to investigate whether silencing multiple immunosuppressive genes in the TME via AAV delivery of Cas13d and gRNAs could function as a combinatorial immunotherapy. We termed this approach MUCIG (<u>M</u>ultiplex <u>U</u>niversal <u>C</u>ombinatorial Immunotherapy via <u>G</u>ene-silencing). We first designed different scales of gene library pools targeting combinations of immunosuppressive genes (**Figure 1B**). Our first gene library pool was designed on the basis of several criteria. By leveraging the knowledge from the literature and the immunogenomic databases such as TISIDB ^24^, we identified 588 tumor immunosuppressive genes and 535 tumor immunostimulatory genes (**Figure 1B**). In order to avoid undesired side effects, we excluded these tumor immunostimulatory genes. We also excluded tumor suppressor genes (TSGs) to avoid potential pro-tumor effect by TSG knockdown, and excluded house-keeping genes to avoid potential toxicity associated with killing normal cells by essential gene knockdown. We further considered the top hits identified from functional screens for genetic factors that enable cancer cells to escape the immune system ^25–28^, selecting genes that have been experimentally validated to be cancer immunotherapy targets. Next, we identified a core set of genes which were recently identified as cancer-intrinsic T cell killing evasion genes across at least 3 cancer models ^29^. Thus, we curated a total of 125 genes from screen data. With a tiered approach, we designed four initial Cas13d gRNA pools for MUCIG experiments (MUCIG-pool1: 313 genes, pool2: 152 genes; pool3: 55 genes; pool4: 19 genes) (**Figure 1B**). We designed Cas13d gRNA pools targeting these gene pools, with 5 gRNAs per gene for most genes.

To facilitate direct delivery of these pools into tumors, we generated an all-in-one AAV vector (AAV-U6-gRNAs-EFS-Cas13d) (**Figure 1A**), which includes both Cas13d and guide-RNA. We synthesized and cloned the gRNA pools and produced the four AAV-MUCIG viral pools accordingly. To evaluate the *in vivo* efficacy of these gene pools against tumors, we first utilized a syngeneic orthotopic tumor model of triple negative breast cancer (TNBC) (E0771 in C57BL/6 mice), which is known to be moderately responsive to immunotherapy. C57BL/6Nr (B6) mice bearing E0771 fat pad transplanted tumors were treated with AAV-MUCIG pools by intratumoral viral administration (**Figure 1C**). All AAV-MUCIG-pools treatment led to significantly reduced tumor burden compared to the AAV-vector or PBS treatment (**Figure 1D**). These data indicated that all four compositions of AAV-MUCIG treatment had therapeutic effect in this tumor model. Different scale and composition of gene pools showed different extent tumor burden reduction effect, among the 4 gene pools, MUCIG-pool2 and 4 showed better therapeutic effect than pool1 and pool3 (**Figure 1D**). This could be influenced by the composition of the genes, the relative concentration of each gRNA, the effects of silencing, and other combinatorial effects.

### A four-gene AAV-MUCIG composition elicits potent anti-tumor immunity

While the AAV-MUCIG gene pools had evidence of anti-tumor responses, we reasoned that further optimization and simplification of the library might increase treatment efficacy by reducing the proportion of potential neutral or detrimental gRNAs that are delivered to the tumor. To further refine the MUCIG-pool, but still aim to find a universal functional combinations across a variety of tumor types, we assessed protein-level expression of the genes targeted in MUCIG-pool4 across a panel of syngeneic cancer cell lines. We excluded the genes that are primarily expressed in non-tumor cells, such as the T cell checkpoints *Pdcd1, Lag3*, and *Havcr2/Tim3*. In addition to the genes targeted in pool4, we also tested other known immunosuppressive genes, such as *Tgf-β*. We systematically analyzed 17 genes by flow cytometry, both for surface and intracellular expression, in ten different syngeneic cell lines across nine different cancer types (MB49 bladder cancer, MC38 colon cancer, Hepa liver cancer, GL261 brain cancer, Pan02 pancreatic cancer, A20 lymphoma, Colon 26 colon cancer, E0771 breast cancer, B16F10 melanoma, and LLC lung cancer lines) (**Figure 2A, B**). Through this unbiased combined immune gene expression analysis, we pinpointed 4 genes (*Pdl1/Cd247, Cd47, Galectin9/Lgals9,* and *Galectin3/Lgals3*) that were abundantly expressed at the protein-level across different cancer cell types. We also examined the human cancer gene expression database and confirmed that the human orthologs of these genes are expressed across a variety of human tumors, supporting their clinical relevance ^30^.

**Figure 2.**
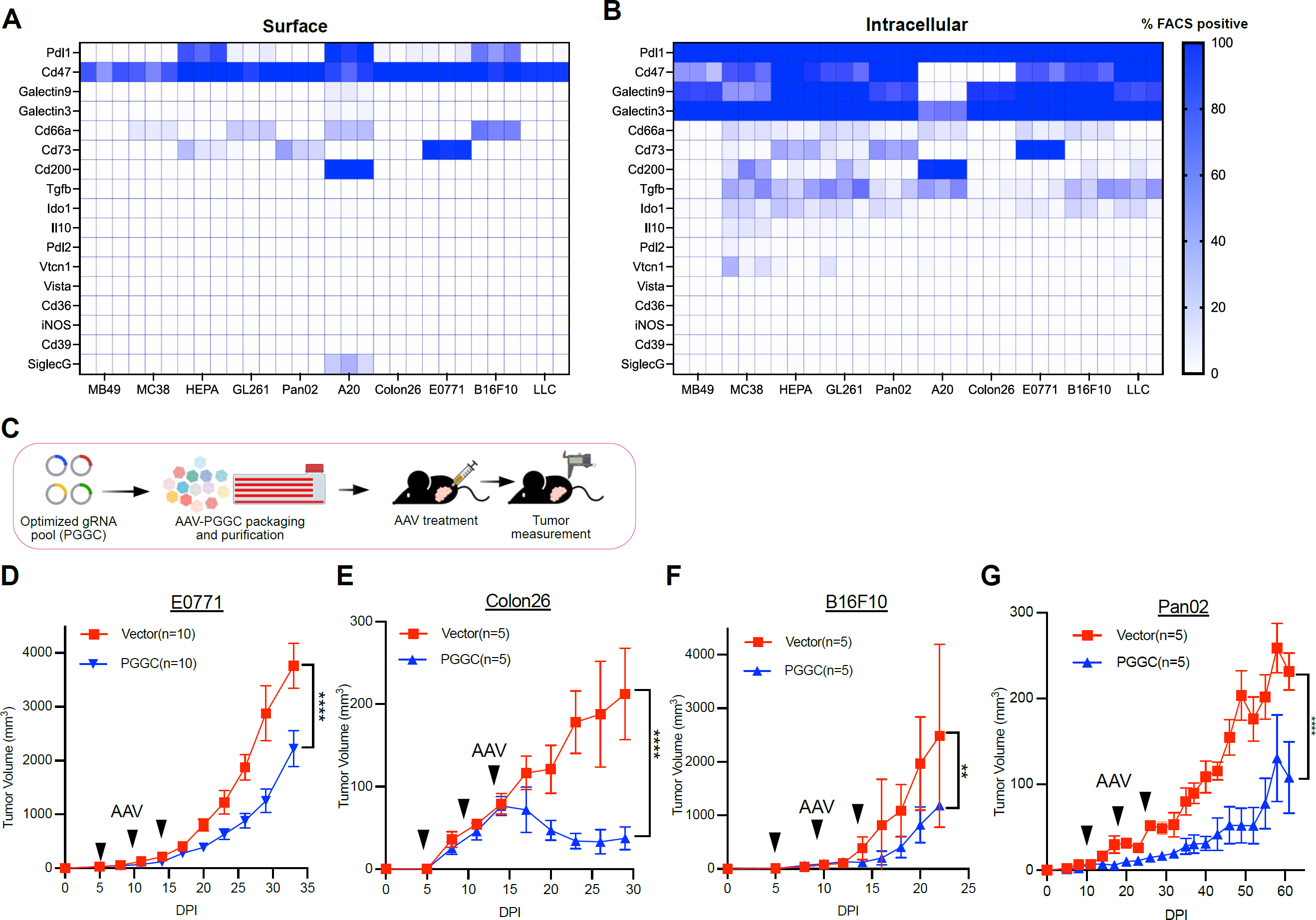
Rational optimization of MUCIG generates AAV-PGGC and demonstrates broader anti-tumor activity in syngeneic models of different cancer types. **A-B.** Protein-level characterization of a list of immunosuppressive factors across a panel of syngeneic cancer cell lines. Heat maps detailing surface (A) and intracellular (B) expression of all assayed immunosuppressive factors determined by flow cytometry. Data are expressed in terms of the percentage of total cells that express each marker. **C.** Schematics of the experimental design for intratumoral delivery of the four-gene AAV-PGGC cocktail. Rational optimization of MUCIG generates AAV-PGGC, an effective four-gene combination immunotherapy, PGGC (PDL1, GALECTIN9, GALECTIN3, and CD47) **D-G.** Growth curves with AAV-PGGC treatment in 4 tumor models. Mice were intratumorally injected with AAV-Vector (Cas13d) and AAV-PGGC) at the timepoints indicated by black arrowheads with 2e^11^ AAV per dose. (D) Breast cancer, (E) Colon cancer, (F) Melanoma model, and (G) Pancreatic cancer. Data points in this figure are presented as mean ± s.e.m. Statistical significance was assessed by two-way ANOVA. ** *p* < 0.01, **** p < 0.0001. Non-significant comparisons not shown.

From the flow data, GALECTIN9 and GALECTIN3 were exclusively expressed intracellularly among all cell lines (**Figure S4A**). Of note, current standard monoclonal antibodies can not inhibit such intracellular targets, however this is achievable by Cas13d-mediated silencing. CD47 was highly expressed on the surface and also expressed intracellularly (**Figure S4A**). Surprisingly, PDL1 was highly expressed intracellularly, even in cell lines with absent surface expression of PDL1 (**Figure S4A**). Since immune checkpoints are often induced in the process of tumorigenesis, we tested expression of these genes in an *in vivo* E0771 tumor model, by flow cytometry analysis of these four proteins in dissociated single cells from tumor samples. Our results showed that all four factors (PDL1, CD47, GALECTIN9 and GALECTIN3) were expressed in both tumor and immune cells (**Figure S4B**).

We then designed a gRNA composition targeting these four genes as a rational and simplified version of MUCIG (named **PGGC** for ***P****dl1;**G**alectin9;**G**alectin3;and **C**d47*), with one of the top gRNA for each gene. We then delivered the AAV-PGGC pool into tumor-bearing mice by intratumoral injection (**Figure 2C**). We found that treatment with AAV-PGGC led to significant reduction of tumor growth in 4 representative models with different levels of responsiveness to immune checkpoint blockade antibody therapeutics, including E0771 breast cancer (moderate) (**Figure 2D**), Colon26 colon cancer (sensitive) (**Figure 2E**), B16F10 melanoma (resistant) (**Figure 2F**), and Pan02 pancreatic cancer (resistant) (**Figure 2G**) mouse models. These data suggest that this four gene formula of Cas13d/gRNA-pool (AAV-PGGC) is effective across different cancer types in animal models.

### AAV-PGGC treatment promotes T cell tumor infiltration while hampering the recruitment of immunosuppressive cells

We then sought to examine how AAV-PGGC treatment influence the immune composition of the TME. By flow cytometry analysis, we profiled tumor-infiltrating lymphoid and myeloid cell populations in mice that received either PBS, AAV-vector, or AAV-PGGC treatment in two different syngeneic tumor models (E0771 and Colon26) (**Figure 3A, Figure S5**). In the E0771 tumor model, we observed significantly more CD45^+^ tumor infiltrating immune cells in the AAV-PGGC treated mice than the Vector control group (**Figure 3B**). Among the tumor infiltrating lymphocytes (TILs), we also found a significant increase of CD8^+^ and CD4^+^ T cells in the AAV-PGGC treated mice compared to Cas13d-vector control (**Figure 3B**). In addition, though there were no substantial changes in the macrophage or the DC population, between AAV-PGGC and Vector control, there was a significant decrease of MDSCs, a heterogeneous cell population with the capacity to functionally suppress T cell responses (**Figure 3B**). In an independent tumor model, Colon26, we similarly observed a significant increase of significantly more CD8^+^ TILs (but not CD4^+^ TILs) in the AAV-PGGC treatment group compared to Vector control group (**Figure S6A**). For the innate populations in the Colon26 model, in AAV-PGGC treated tumors compared to Vector control, there were more tumor-infiltrating macrophages and DCs, relevant to antigen presentation and for priming adaptive immune responses (**Figure S6A**). In AAV-PGGC treated tumors compared to PBS control, again there was a significant decrease of MDSCs (**Figure S6A**).

**Figure 3.**
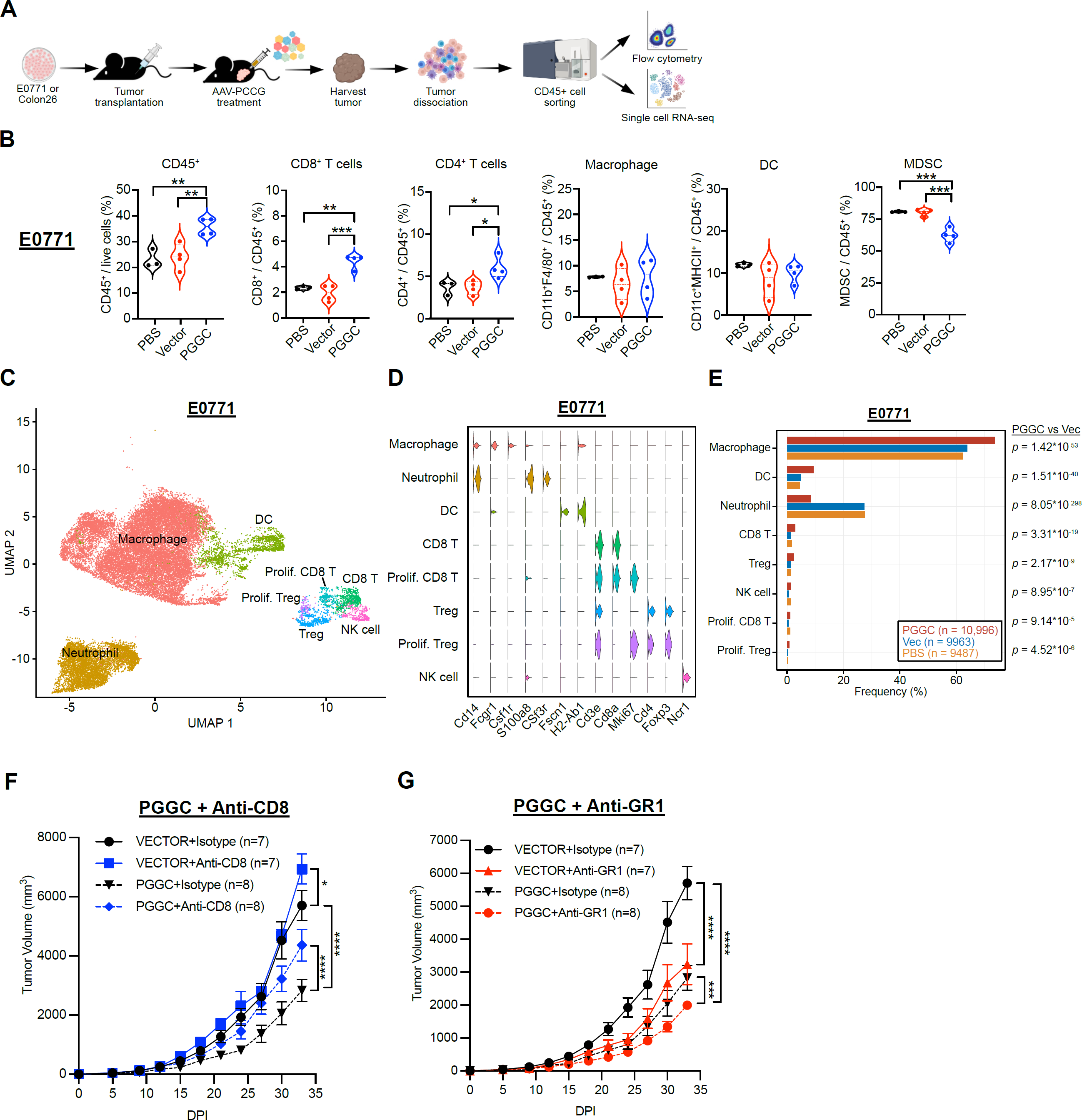
AAV-PGGC treatment remodels the immunosuppressive tumor microenvironment and inhibits metastatic cancer combined with Anti-GR1. **A.** Schematic of experimental design for analyzing of the composition of tumor infiltrating immune populations after AAV-PGGC therapy. **B.** Relative abundances of several immune populations in orthotopic E0771 tumors, at the endpoint of tumor study (35 days post tumor induction). For the E0771 model, mice were intratumorally treated with PBS (n = 3), AAV-Vector (n = 4) or AAV-PGGC (n = 4) at days 4, 9 and 14. Statistical significance was assessed by one-way ANOVA Tukey’s multiple comparisons test, adjusted P Value. (* *p* < 0.05, ** *p* < 0.01, *** *p* < 0.001). Non-significant comparisons not shown. **C.** UMAP visualization of single tumor-infiltrating immune cells, profiled by scRNA-seq. Mice bearing orthotopic E0771 tumors were treated with PBS, AAV-Vector or AAV-PGGC at days 4, 9 and 14. Tumors were harvested at day 29, and live CD45^+^ cells were sorted for scRNA-seq. **D.** Violin plots showing the expression levels of representative marker genes across the main cell clusters. **E.** Relative proportions of each cell type, across treatment groups. Statistical analysis between groups was performed by two-tailed Fisher’s exact test. **F-G.** Growth curves of AAV-PGGC plus antibody combination treatment in syngeneic orthotopic model (E0771 in C57BL/6 mice). (F) PGGC plus anti-CD8 against E0771 breast tumors in C57BL/6, (G) PGGC plus anti-GR1 against E0771 breast tumors in C57BL/6. Data points in this figure are presented as mean ± s.e.m. Statistical significance was assessed by two-way ANOVA, or Log-rank test as indicated in each panel. * *p* < 0.05, ** *p* < 0.01, *** *p* < 0.001, **** p < 0.0001. Non-significant comparisons not shown.

To systematically investigate the effect of AAV-PGGC treatment on the immune cell populations and their transcriptomics in the TME, we performed single-cell RNA-seq (scRNA-seq) of tumor-infiltrating immune cells in mice treated with PBS, AAV-Vector, or AAV-PGGC (**Figure 3C,D**). Consistent with the flow cytometry analysis, scRNA-seq of the E0771 tumor model revealed significant changes in multiple immune cell populations after AAV-PGGC treatment (**Figure 3E**), including an increase of CD8^+^ T cells and proliferating CD8^+^ T cells. Similarly, in the Colon26 model, we observed more CD8^+^ T cells and proliferating CD8+ T cells with AAV-PGGC treatment (**Figure S6B,C**). On the other hand, there was a substantial reduction of neutrophil abundances in AAV-PGGC treatment group compared with PBS or vector control group (**Figure 3E**), which was also observed in the Colon26 model (**Figure S6D**).

Via differential expression analysis (DE) of sc-RNA-seq data, we identified DE genes in the cell types whose abundances were most affected by AAV-PGGC, including CD8^+^ T cells, neutrophils and macrophages. We found a panel of genes associated with key immunosuppressive functions were downregulated across both E0771 and Colon26 models, including *Arg2, Il1b, Trem1, S100a8, S100a9, Tigit,* and *Cd37* (**Figure S6E-G**). It was reported that CD37 could inhibit Cd3-induced T cell proliferation^31^. In CD8^+^ T cells, we found *Cd37* was downregulated in AAV-PGGC treatment group when compared with vector group (**Figure S6E**). *Tigit*, a marker of T cell exhaustion ^32^, was also decreased in CD8^+^ T cells of AAV-PGGC treatment group (**Figure S6E**). *Arg2*, which has been implicated in the immunosuppressive functions of neutrophils, was downregulated in the AAV-PGGC group along with *Ifitm1* and *Ifitm3*, two genes that play a role in suppressing interferon mediated immunity ^33, 34^ (**Figure S6F**). *S100a8* and *S100a9*, two factors that help recruit MDSCs to the TME ^35^, were downregulated in macrophages and CD8^+^ T cells from AAV-PGGC treated tumors (**Figure S6E,G**). Consistent with the observed reductions in tumor-associated neutrophils after AAV-PGGC treatment, the genes encoding neutrophil-recruiting chemokines *Cxcl1* and *Cxcl2* were significantly downregulated in both neutrophils and macrophages isolated from tumors treated with AAV-PGGC (**Figure S6F,G**). These data suggested that AAV-PGGC treatment can effectively reverse the immunosuppressive TME, promoting T cell infiltration and reducing suppressive myeloid cell populations.

### AAV-PGGC combined with anti-GR1 treatment further inhibit tumor growth

Given the increase of CD8^+^ T cells and reduction of neutrophils in the TME after AAV-PGGC treatment, we next tested how these two cell populations influence the therapeutic efficacy of AAV-PGGC. We performed CD8^+^ T cell or MDSC/neutrophils depletion by *in vivo* injection of anti-CD8 or anti-GR1 antibody, respectively (**Figure 3F**). We observed that mice with CD8^+^ T cell depletion partially impaired the anti-tumor effect of AAV-PGGC (**Figure 3F**), which indicate that AAV-PGGC treatment is partially dependent on CD8^+^ T cells. Because the previous TIL analysis indicated large amount of myeloid derived immunosuppressive cells (MDSCs) even after AAV-PGGC treatment (**Figure 3B,C**), we wondered whether the combined treatment with anti-GR1 to deplete MDSCs could enhance the therapeutic efficacy. Results showed that depletion of MDSCs and neutrophils by anti-GR1 in combination with AAV-PGGC treatment could further reduce the tumor burden when comparing to either AAV-PGGC or anti-GR1 antibody alone, again in a syngenetic orthotopic E0771 breast cancer model in C57BL/6 mice (**Figure 3G**). These data suggested that CD8^+^ T cells and MDSCs/neutrophils together play critical roles in AAV-PGGC therapy, where depletion of CD8 T cells by anti-CD8 reduced PGGC’s therapeutic efficacy, and combinatorial treatment with anti-GR1 enhanced PGGC’s efficacy.

The TME is enriched with immunosuppressive factors that can be derived from tumor cells, tumor-associated fibroblasts or the infiltrating immunosuppressive cells ^36–38^. A number of preclinical studies have demonstrated that neutralization of immunosuppressive factors can reverse the immunosuppressive TME and promote anti-tumor immunity ^39, 40^. Various strategies have been developed to repress such targets or their activity, including siRNAs, antisense oligos, antagonistic antibodies, and small molecule inhibitors. Here we take an approach by simultaneously repressing multiple immunosuppressive genes directly in the TME. We leverage the modularity of the CRISPR/Cas13d system to devise multiple combinatorial immunotherapies, demonstrating the anti-tumor efficacy of several different libraries of varying complexity. Because multiplexing gRNAs is simple, it is readily feasible to generate gRNAs pools that target a custom number of immunosuppressive genes on demand.

Key challenges with tumor gene therapy include on-target-specificity and gene delivery efficiency. Cas13d binds and cleaves single-strand RNA, thus avoiding safety concerns stemming from unintended DNA damage caused by Cas9 or Cas12a. In addition, Cas13d is more compact compared to Cas9, Cas12a, and many other Cas13 family members, conferring a key advantage for viral vector delivery ^16^. We utilized AAVs to deliver the Cas13d-gRNA payload into tumors, as AAVs can efficiently deliver foreign genetic materials *in vivo* with minimal toxicity. Indeed, we observed persistent exogenous gene expression up to two weeks after the final intratumoral injection of AAV. However, as we observed here, one potential safety limitation of intratumoral AAV delivery is the propensity for AAVs to transduce cells in the liver, although we did not observe obvious gross side effects in any of the MUCIG-treated mice. AAV-mediated delivery still poses safety concerns relative to non-viral approaches, as the AAV genome can integrate into the host cell genome and double-strand break sites ^41^. In addition, the diversity of immunosuppressive pathways that are engaged across different tumors poses an important challenge. Nevertheless, the MUCIG approach, with the versatility of targeting virtually any reasonable combinations of genes using CRISPR-Cas13d and gRNA pools, offers far greater flexibility and modularity compared to conventional antagonistic antibodies or small molecules. By further customizing the cocktail of immunosuppressive factors that is targeted by MUCIG, or by utilizing more specific delivery vehicles, we anticipate that the therapeutic window can be optimized to minimize off-tumor toxicity while maintaining anti-tumor efficacy.

In summary, here we present MUCIG, a versatile strategy for combinatorial cancer immunotherapy by multiplexed targeting of the immunosuppressive gene collections. Taking advantage of the multiplex capability of Cas13d, MUCIG can simultaneously inhibit multiple immune checkpoints tailored to specific cancer types or commonly across multiple cancer types, simply by designing and putting together gRNA combinations. Using tissue-specific promoter or other control elements, MUCIG’s application can be extended to tissue-targeted multiplexed gene silencing beyond oncology. MUCIG can be rapidly customized for targeting any combinations of genes in diverse tissue or cell types *in vivo*.

## Author Contributions

Conceptualization: SC. Design: FZ, GW, RC, SC. Experiment lead: FZ, GW. Analytic lead: RC. Experiment assistance and support: EH, MM, YZ. Manuscript prep: FZ, RC, GW, SC. Supervision and funding: SC.

## Acknowledgments

### Institutional approval

This study has received institutional regulatory approval. All recombinant DNA and biosafety work was performed under the guidelines of Yale Environment, Health and Safety (EHS) Committee with an approved protocol (Chen-rDNA-15-45). All animal work was approved by Yale University’s Institutional Animal Care and Use Committee (IACUC) and performed with approved protocols (Chen #2018-20068; #2021-20068).

### Discussion and Support

We thank all members in Chen laboratory, as well as various colleagues in Yale Genetics, SBI, CSBC, MCGD, Immunobiology, BBS, YCC, YSCC, and CBDS for assistance and/or discussions. We thank various Yale Core Facilities such as YCGA, HPC, WCAC, KBRL for technical support.

### Funding

S.C. is supported by NIH/NCI/NIDA (DP2CA238295, R01CA231112, R33CA225498, RF1DA048811), DoD (W81XWH-17-1-0235, W81XWH-20-1-0072, W81XWH-21-1-0514), Damon Runyon Dale Frey Award (DFS-13-15), Melanoma Research Alliance (412806, 16-003524), Cancer Research Institute (CLIP), AACR (17-20-01-CHEN), The V Foundation (V2017-022), Alliance for Cancer Gene Therapy, Sontag Foundation (DSA), Pershing Square Sohn Cancer Research Alliance, Dexter Lu, Ludwig Family Foundation, Blavatnik Family Foundation, and Chenevert Family Foundation. GW is supported by CRI Irvington and RJ Anderson Postdoctoral Fellowships. RC is supported by NIH MSTP training grant (T32GM007205) and NRSA fellowship (F30CA250249).

### Data and material availability

All data generated or analyzed during this study are included in this article and its supplementary information files. Source data and statistics are provided in an excel file of **Source data and statistics**. Processed data for genomic sequencing and gene expression are provided as processed quantifications in

### Supplementary Datasets

Genomic sequencing raw data are being deposited to NIH Sequence Read Archive (SRA) and/or Gene Expression Omnibus (GEO), with pending accession numbers. Data, codes and materials that support the findings of this research are available from the corresponding author upon reasonable request to the academic community.

## Methods

### Cell lines

HEK293FT cell was purchased from ThermoFisher Scientific for producing viruses. All cell lines used in this paper were maintained at 37C with 5% CO2 in D10 medium (Dulbecco’s modified Eagle’s medium supplemented with 10% fetal bovine serum).

### Mice

Mice of both sexes, between age 6 and 12 weeks, were used for the study. 6-8-week-old C57BL/6Nr mice were purchase from Charles River lab. Female mice were used for breast cancer (E0771) models. Male mice were used for B16F10 and Pan02 mouse model. 6-8-week-old BALB/C mice were purchased from Jackson lab, which were used for Colon26 mouse model. All animals were housed in standard, individually ventilated, pathogen-free conditions, with a 12 h:12 h or a 13 h:11 h light cycle, at room temperature (21– 23 °C) and 40–60% relative humidity.

### Cas13d cancer cell line generation

For lentivirus production, 20µg plasmid of PXR001 (EF1a-Cas13d-2A-EGFP, addgene#109049) together with 10µg pMD2.G and 15µg psPAX2 were co-transfected into HEK293FT cells in a 150mm cell culture dish at 80-90% confluency using 135µl LipoD293 transfection reagent (Signage, SL100668). Virus supernatant was collected 48h post transfection, centrifuged at 3000g for 15min to remove the cell debris. The supernatant was then concentrated with Amicon Ultra-15 filter from 20ml to 2ml. The virus was aliquoted and stored at -80C. To generate Cas13d overexpression cell line, the cancer cells were transduced with lentivirus PXR001, and the positive cells which were GFP expressing were flow cytometry sorted.

### Transfection and flow cytometry knockdown efficacy test

To test each gRNA knockdown efficacy, gRNAs were cloned into BbsI site of PXR003 plasmid (Cas13d gRNA cloning backbone, addgene#109053) and were transient transfected into Cas13d expressing cancer cell. For the transfection experiments, 5×10^4^ cells per well of a 48 well plate was seeded 12h before transfection. 500ng gRNA plasmid together with a 1:1 ratio of Lipofectamine 2000 to DNA were transfected into cells. Flow cytometry was performed at 48h post transfection.

### Dual reporter cell line with Cas13d expressing generation

A lentivirus version plasmid expressing Cas13d and blasticidin (EF1a-Cas13d-T2A-BSD-WPRE) was cloned. A lentivirus version plasmid expressing U6_Direct repeats_guideRNA was cloned. E0711 cell line was co-transduced with three lentiviruses (Cas13d-blasticidin, GFP and mCherry). The Cas13d-expressing dual reporter E0771 cells was selected with blasticidin and then sorted with GFP^+^ mCherry^+^ double positive cells. The dual reporter cells were then transduced with Cas13d-guideRNA lentivirus.

### Quantitative reverse transcription PCR (qRT-PCR)

RNA was extracted by TRIzol (Invitrogen), and the cDNA was synthesized using the PrimeScript RT master kit (Takara, RR036A). The qPCR was done using PowerUp SYBR Green Master Mix (Thermofisher) following the instruction. The expression levels of genes were detected on QuantStudio™ 3 Real-Time PCR System. The gene relative expression was calculated by the 2-ΔΔCt method. GAPDH was measured as reference.

### Generation of AAV-MUCIG pools

An AAV version plasmid expressing U6-mutation direct repeat-gRNA clone site-EFS-Cas13d (pAAV-U6-EFS-Cas13d) was cloned into AAV backbone. All pooled gRNA library were synthesized as single stranded oligonucleotides from Genescript or IDT. The oligos were amplified by PCR and Gibson cloned into pAAV-U6-EFS-Cas13d. The purification and electroporation of Gibson products into Endura electrocompetent cells were performed as previously described^42^, with at least x100 coverage of colonies represented per sgRNAs. AAV was produced by co-transfecting HEK293FT cells with AAV-MUCIG pool together with AAV9 serotype plasmid and helper plasmid PDF6. Briefly, HEK293FT cells were seeded in 150cm dish or hyper flask 12-18h before transfection. When cells got 80-90% confluency, 6.2µg AAV-vector or AAV-MUCIG pool, 8.7µg AAV9 serotype, and 10.4µg PDF6 were transfected with 130µl PEI, incubating 10-15min before adding into cells. Replicates collected multiple dishes were pooled to enhance production yield. Cells were collected 72h post transfection. For AAV purification, chloroform (1:10 by volume) was added and was shaken vigorously for 1h at 37°C. NaCl was added to a final concentration of 1M and shaken until dissolved. The mixture was centrifuges at 20,000g for 15min at 4C. The aqueous layer was transferred to a new tube, and then PEG 8000 (10%, w/v) was added and shaken until dissolved. The mixture was incubated on ice for 1h. The pellet was spun down at 20,000g for 15min at 4°C. The supernatant was discarded, and the pellet was resuspended in DPBS. The resuspension was treated with Benzonase and MgCl2 AT 37C for 30min. Chloroform (1:1 by volume) was then added, shaken and spun down at 12,000g for 15min at 4°C. The aqueous layer was isolated and concentrated through Ambion Ultra-15 tube. The concentrated solution was washed with PBS and the filtration process repeated. Then AAV was treated with DNase I for 30min at 37°C. Genomic copy number (GC) of AAV was determined by real-time qPCR using custom TaqMan assays (Thermo Fisher Scientific) targeted to EFS promoter.

### Therapeutic testing of AAV-MUCIG in syngeneic tumor models

Syngeneic orthotopic breast tumor was established by transplanting 2×10^6^ E0771 cells into mammary fat pad of 6–8-week-old female C57BL/6Nr mice. Then 5, 9, and 14 days after transplantation, 2e^11^ AAV partials of vector or MUCIG, or PBS were injected intratumorally into tumor bearing mice. The tumor volume was measured every 3-4 days. For the B16F10 melanoma model, 1×10^6^ B16F10 cancer cells were subcutaneously injected into the male left flank of C57BL/6Nr mice. 5, 9, 13 days post transplantation, 2e^11^ AAV partials of vector or MUCIG, or PBS were intratumorally administrated into tumor bearing mice. The tumor volume was measured every 2 days. For the pancreatic tumor model, 2×10^6^ Pan02 cells were subcutaneously injected into the left flank of C57BL/6Nr mice. Then, 5, 14, 18 days after transplantation, 2e^11^ AAV partials of vector or MUCIG, or PBS were intratumorally administrated into tumor bearing mice. The tumor volume was measured every 3-4 days. For the colon tumor model, 2×10^6^ Colon26 cells were subcutaneously injected into the left flank of BALB/C mice. Then, 5, 9, 14 days after transplantation, 2e^11^ AAV partials of vector or MUCIG, or PBS were intratumorally administrated into tumor bearing mice. The tumor volume was measured every 3 days. Tumor volume was calculated with the formula: volume = π/6*xyz. Two-way ANOVA was used to compare growth curves between treatment groups.

### *In vivo* luciferase imaging

The bioluminescent imaging was performed to detect AAV delivery gene expression. Mice were injected with luciferin (150mg/kg) by intraperitoneal injection and activity quantified in live animal for 10min later following with 1min exposure. The photon flux was monitored by the PE IVIS Spectrum in vivo imaging system. The signaling was monitored and quantified by the IVIS software.

### Isolation of TILs

Tumors were minced into 1 mm size pieces and then digested with 100U/ml collagenase IV and DNase I for 60min at 37°C. Tumor suspensions were filtered through 100-μm cell strainer to remove large bulk masses. The cells were washed twice with wash buffer (PBS plus 2% FBS). 1ml ACK lysis buffer was added to lysis red blood cell by incubating 2-5 min at room temperature. The suspension was then diluted with wash buffer and spin down at 400g for 5min at 4°C. Cell pellet was resuspended with wash buffer and followed by passing through a 40μm cell strainer. Cells were spin down and washed twice with wash buffer. At last, cell pellet was resuspended in MACS buffer (PBS with 0.5% BSA and 2mM EDTA). The single cell suspensions were used for flow cytometry staining and FACS sorting. TILs were labeled as CD45 positive cells.

### Flow cytometry

For the TILs flow cytometry analysis, single cell suspension from tumor were prepared as described above. For the myeloid cell staining panel, anti-CD45-Percp-Cy5.5, anti-CD11b-FITC, anti-CD11c-PE/Dazzle, anti-F4/80-PE, anti-Ly6G-BV605, anti-Ly6C-APC, and anti-MHCII-PE/Cy7 were used. For lymphoid cell staining panel, anti-CD45-Percp-Cy5.5, anti-CD8-BV605, anti-CD4-PE. All flow antibodies were used at 1:100 dilutions for staining. The LIVE/DEAD Near-IR was diluted 1:1000 to distinguish live or dead cells.

For the in vitro cancer cell line staining, cancer cells were incubated with trypsin and washed twice with PBS. For cell surface staining, surface antibody was diluted 1:100 and stained in MACS buffer on ice for 15min. Cells were washed twice with MACS buffer. For intracellular staining, Intracellular Fixation & Permeabilization Buffer Set (eBioscience) was used to fix and permeabilize cells. Briefly, after the surface marker staining, cells were resuspended in 100μl Fixation/Permeabilization working solution, and incubated on ice for 15 min. Then cells were washed with 1× permeabilization buffer by centrifugation at 600g for 5 min. Then the cell pellet was resuspended in 100μl of 1× permeabilization buffer with 1:100 intracellular staining antibodies and incubating on ice for 15min. After staining, cells were centrifuged at 600g for 5 min, and washed twice with staining buffer before being analyzed or sorted on a BD FACSAria. The data were analyzed using FlowJo software.

### Immune cell profiling by scRNA-seq

E0771 or Colon26 tumors were collected at the indicated time point post injection. Single cell suspensions were collected as described above. The cells were labeled with CD45-Percp-Cy5.5 antibody and live/dead dye. FACS sorted cells were gated on CD45^+^ live cells. Sorted cells were washed with PBS, and cell numbers and viabilities were assessed by trypan blue staining. The 10,000 CD45^+^ cells isolated from tumors were used for scRNA-seq library prep by following the protocol from 10× Genomics Chromium Next GEM Single Cell 5’ Reagent Kits V2.

### scRNA-seq data analysis

Analysis of scRNA-seq was performed using the Seurat v4 package in R. All cells from the three treatment groups (PBS, AAV-Vector, and AAV-PGGC) were merged and integrated by tumor type (E0771 or Colon26). The data was filtered to retain cells with < 15% mitochondrial counts and 200-3500 unique expressed features. The expression data for each cell was normalized by the total reads and log-transformed. We utilized Harmony to integrate datasets from the same tumor type for the purpose of identifying cell clusters. Each cell cluster was annotated by cell type using canonical marker genes, with higher-resolution subclustering of the lymphocyte populations. To determine differences in cell type frequencies, we constructed 2×2 contingency tables for each cell type, comparing AAV-Vector and AAV-PGGC treatment groups. A two-tailed Fisher’s exact test was performed on the contingency table for each cell type. Differentially expressed genes were identified by comparing cells from AAV-Vector vs AAV-PGGC treatment groups using the default settings in Seurat, with statistical significance set at adjusted *p* < 0.05.

### Statistical analysis

Standard non-NGS statistical analyses were performed in GraphPad Prism using specific statistical tests where appropriate, as detailed in figure legends. NGS statistical analyses were performed in R/RStudio. Different levels of statistical significance were accessed based on specific *p* values and type I error cutoffs (e.g. 0.05, 0.01, 0.001, 0.0001).

## Reporting summaries

### Statistics

For all statistical analyses, we confirmed that the items mentioned in NPG reporting summary are present in the figure legend, table legend, main text, or Methods section.

### Standard statistical analysis

All statistical methods are described in figure legends, methods and/or supplementary Excel tables. Source data and statistics were provided in a supplemental excel table.

### Software and code

#### Data collection

Default softwares in the data collection instruments were used, including Attune focusing cytometer (Attune NxT Software v3.1); FACS Aria Dava; RT-PCR Quant studio.

All deep sequencing data were collected using Illumina Sequencers at Yale Center for Genome Analysis (YCGA).

### Data analysis

Flow cytometry data were analyzed by FlowJo v.10.7.

All simple statistical analyses were done with Prism 9.

All NGS analyses were performed using custom codes.

### Data and resource availability

All data generated or analyzed during this study are included in this article and its supplementary information files. Specifically, source data and statistics for regular experiments are provided in an excel file of Source data and statistics. NGS data are being deposited to SRA/GEO with pending accession codes. Other materials and data are available either through public repositories, or via reasonable requests to the corresponding authors to the academic community.

### Code availability

Custom codes are available either through public repositories, or via reasonable requests to the corresponding author to the academic community.

## Life sciences study design

### Sample size determination

For most cases, each group has five biologically independent samples unless otherwise noted. Details on sample size for each experiment were indicated in methods and figure legends. Sample size was determined according to the lab’s prior work or similar studies in the field.

### Data exclusions

No other data were excluded.

### Replication

Number of biological replicates (usually n >= 3) are indicated in the figure legends. Key findings (non-NGS) were replicated in at least two independent experiments. NGS experiments were performed with biological replications as indicated in the manuscript.

### Randomization

Regular in vitro experiments were not randomized or blinded. Mouse experiments were randomized by using littermates, and blinded using generic cage barcodes and eartags where applicable.

### Blinding

Investigators were blinded to the identity and treatment groups of animals when measuring tumor burden. Investigators were not blinded in *in vitro* experiments. In certain NGS data analysis, investigators were blinded for initial processing of the original data using key-coded metadata.

## Reporting for specific materials, systems and methods

### Antibodies used

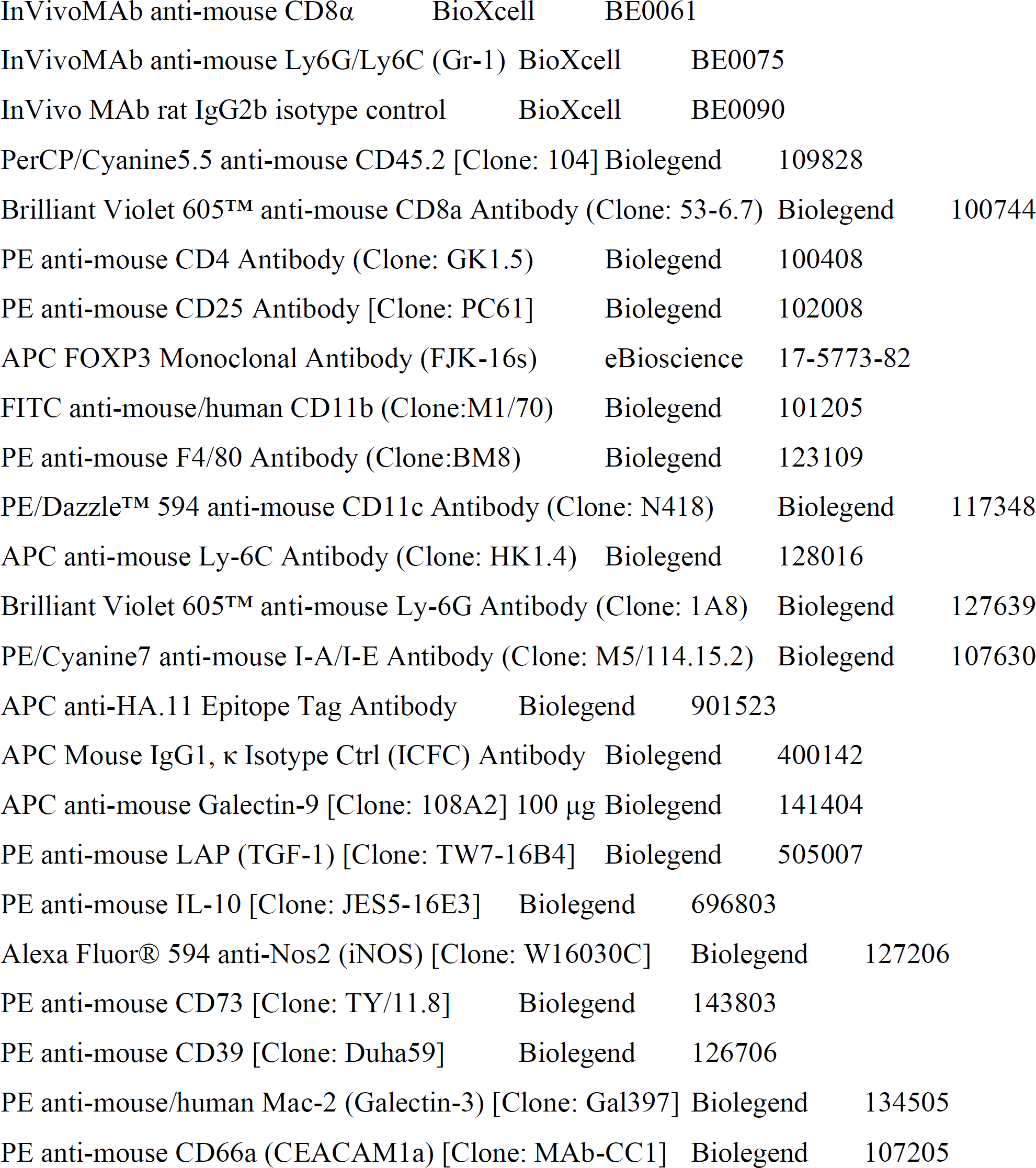
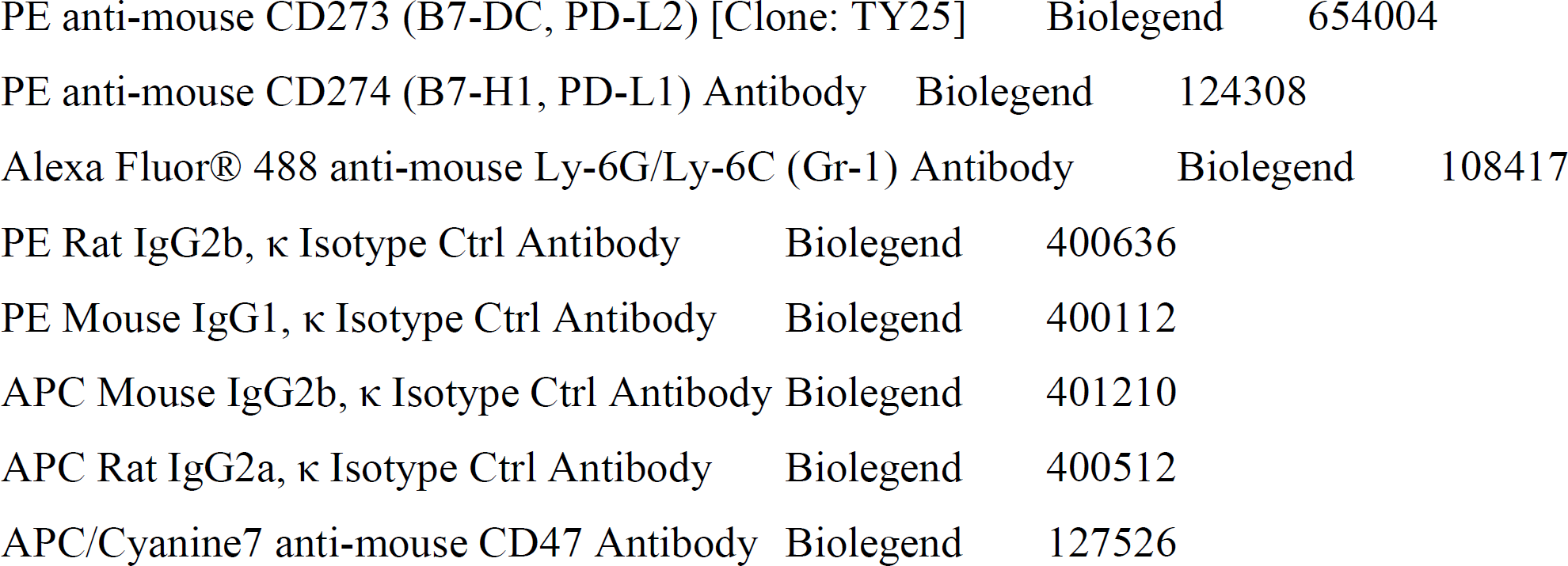

### Antibody validation

Commercial antibodies were validated by the vendors, and re-validated in house as appropriate through antigen-specific experiments.

### Eukaryotic cell lines

Commercial cell lines were acquired from commercial vendors.

HEK293FT ThermoFisher Catalog Number: R70007

E0771 CH3 Catalog Number: 940001

B16F10 ATCC Catalog Number: CRL-6475™

GL261 ATCC Catalog Number: HB-12317™

Pan02 ATCC Catalog Number: CRL-2553™

CT26 ATCC Catalog Number: CRL-2638™

### Authentication

Cell lines were authenticated by original vendors, and re-validated in lab as appropriate by morphology.

### Mycoplasma contamination

All the cell lines used here tested negative for mycoplasma contamination.

### Commonly misidentified lines (See ICLAC register)

No misidentified cell lines were used in the study.

### Animals and other organisms

#### Laboratory animals

C57BL/6Ncr (B6) mice, female 8-12 week mice purchased from Charles River.

#### Wild animals

N/A

#### Field-collected samples

N/A

#### Ethics oversight

The work described in this study is performed under relevant approved protocols (IACUC, rDNA/EHS) in place.

### Flow Cytometry

#### Plots

Confirm the checkboxes of requirements.

### Methodology

#### Sample preparation

Various sample prep details are provided in the Methods section.

#### Instrument

Flow cytometric analysis was performed on an BD FACSAria II or thermo Attune™ NxT.

#### Software

FlowJo v.10.7.1 was used for flow cytometry data analysis.

#### Cell population abundance

N/A.

#### Gating strategy

A lymphocyte gate was defined first from FSC-A v SSC-A. Singlet gates were then defined on FSC-H v FSC-W. Additional gating was performed as described in figure and extended data legends for individual experiments.

#### Gating example figure

Confirmed that a figure exemplifying the gating strategy is provided in the Supplementary Information.

## Supplemental Figure Legends

**Figure S1.**
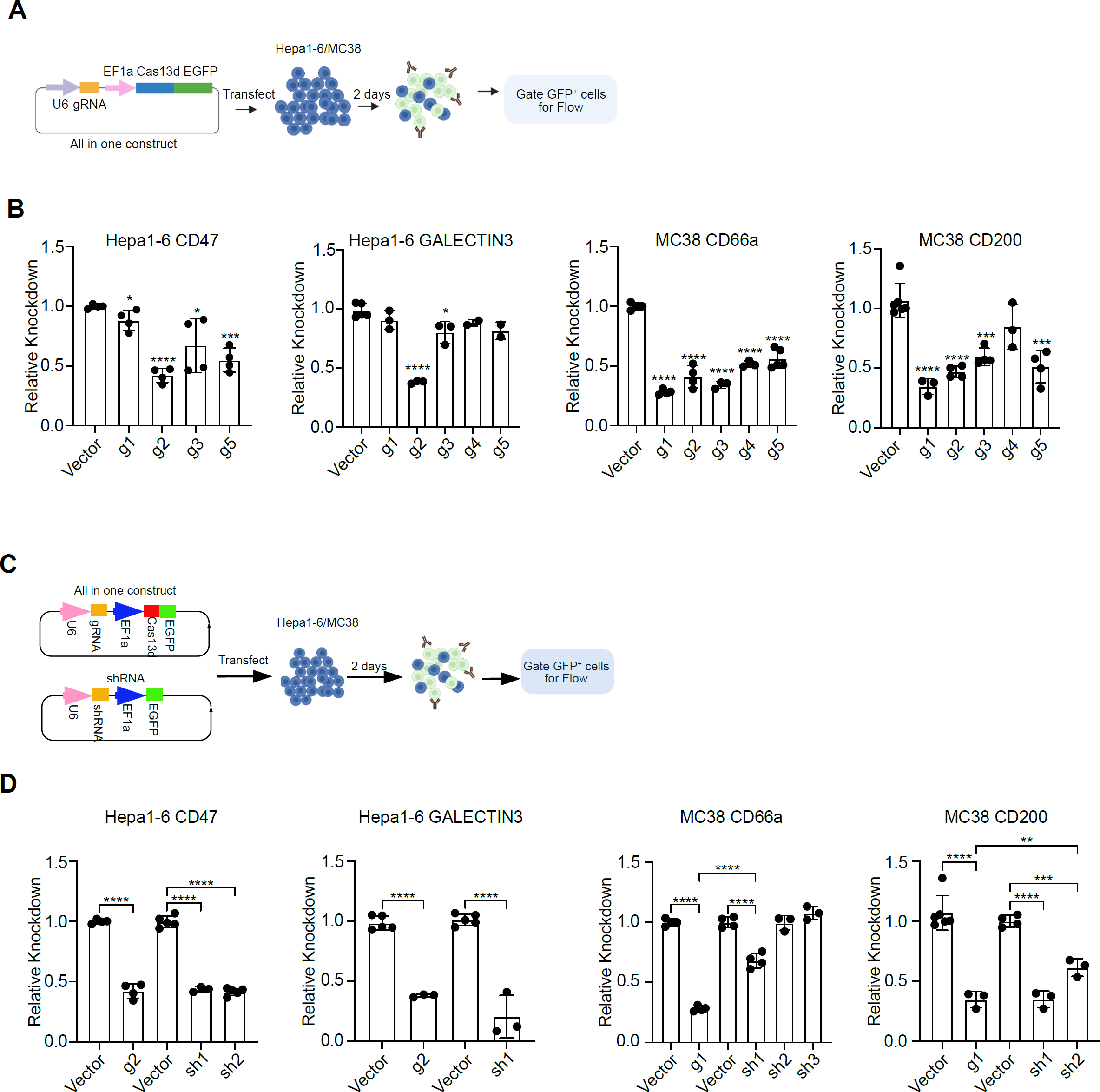
Cas13d-mediated silencing of endogenous immunosuppressive genes in cancer cells. **A.** Schematic of the experimental approach to identify efficient Cas13d gRNAs targeting various immunosuppressive genes. The all-in-one plasmid including gRNA, Cas13 and EGFP was transfected into Hepa1-6 or MC38 cells. Two days after transfection, target gene expression was tested by flow cytometry analysis. The gRNA successful transfected cells were gated by GFP^+^ cells. **B.** Knockdown efficiency of gRNAs targeting different immunosuppressive genes in cancer cell lines. CD47 and GALECTIN3 were tested in Hepa1-6 cells, and CD66a and CD200 were tested in MC38 cells. Data are expressed as the relative mean of fluorescent intensity (MFI). The gene expression level of vector group was normalized to 1. **C.** Schematic of the experimental approach to compare knockdown efficient between Cas13d gRNAs and shRNA. The cas13d all-in-one (gRNA-Cas13-EGFP) or shRNA plasmid was transfected into Hepa1-6 or MC38 cells. Two days after transfection, target gene expression was tested by flow cytometry analysis. The gRNA successful transfected cells were gated by GFP^+^ cells. **D.** Comparison of the Cas13d gRNA-mediated and shRNA-mediated target knockdown. CD47 and GALECTIN3 were tested in Hepa1-6 cells, and CD66a and CD200 were tested in MC38 cells. Data are expressed as the relative mean of fluorescent intensity (MFI). The gene expression level of vector group was normalized to 1. Data points in this figure are presented as mean ± s.e.m. Statistical significance was assessed by two-tailed unpaired *t* test. * *p* < 0.05, ** *p* < 0.01, *** *p* < 0.001, **** p < 0.0001. Non-significant comparisons not shown.

**Figure S2.**
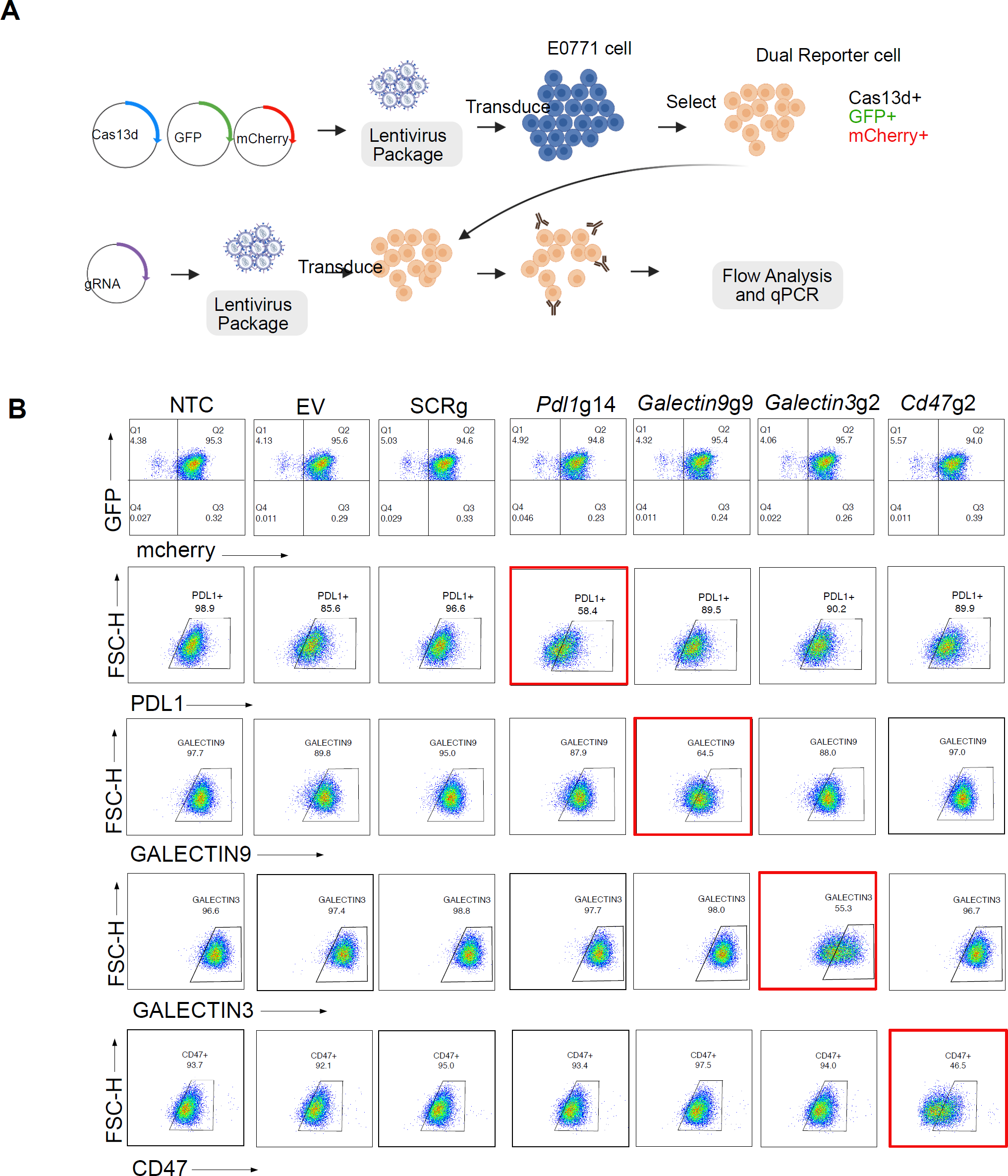
Cas13d on-target and collateral activity testing when targeting endogenous immunosuppressive genes. **A.** Diagram of Cas13d collateral activity and on-target activity by a dual-GFP and mCherry reporter system. E0711 cell line was co-transduced with three lentiviruses (Cas13d-blasticidin, GFP and mCherry). The Cas13d-expressing dual reporter E0771 cells was selected with blasticidin and then sorted by GFP^+^ mCherry^+^ double positive cells. The dual reporter cells were then transduced with Cas13d-guideRNA lentivirus. Then the GFP and mCherry fluorescent signal was determined by flow cytometry. The on-target gene expression was tested by flow cytometry and qPCR. **B.** Flow cytometry analysis of E0771 dual-reporter cells after transduced with different guide RNAs. NTC (Non-Transduced-Control), EV (Empty Vector), SCRg(scramble guideRNA). **C.** RT-qPCR analysis of the target gene expression. The gene mRNA expression level of SCRg was normalized to 1. Data points in this figure are presented as mean ± s.e.m. Statistical significance was assessed by one-way ANOVA Tukey’s multiple comparisons test, adjusted P Value. Multiple comparisons were summarized in the bellowing table. * *p* < 0.05, ** *p* < 0.01, *** *p* < 0.001.

**Figure S3.**
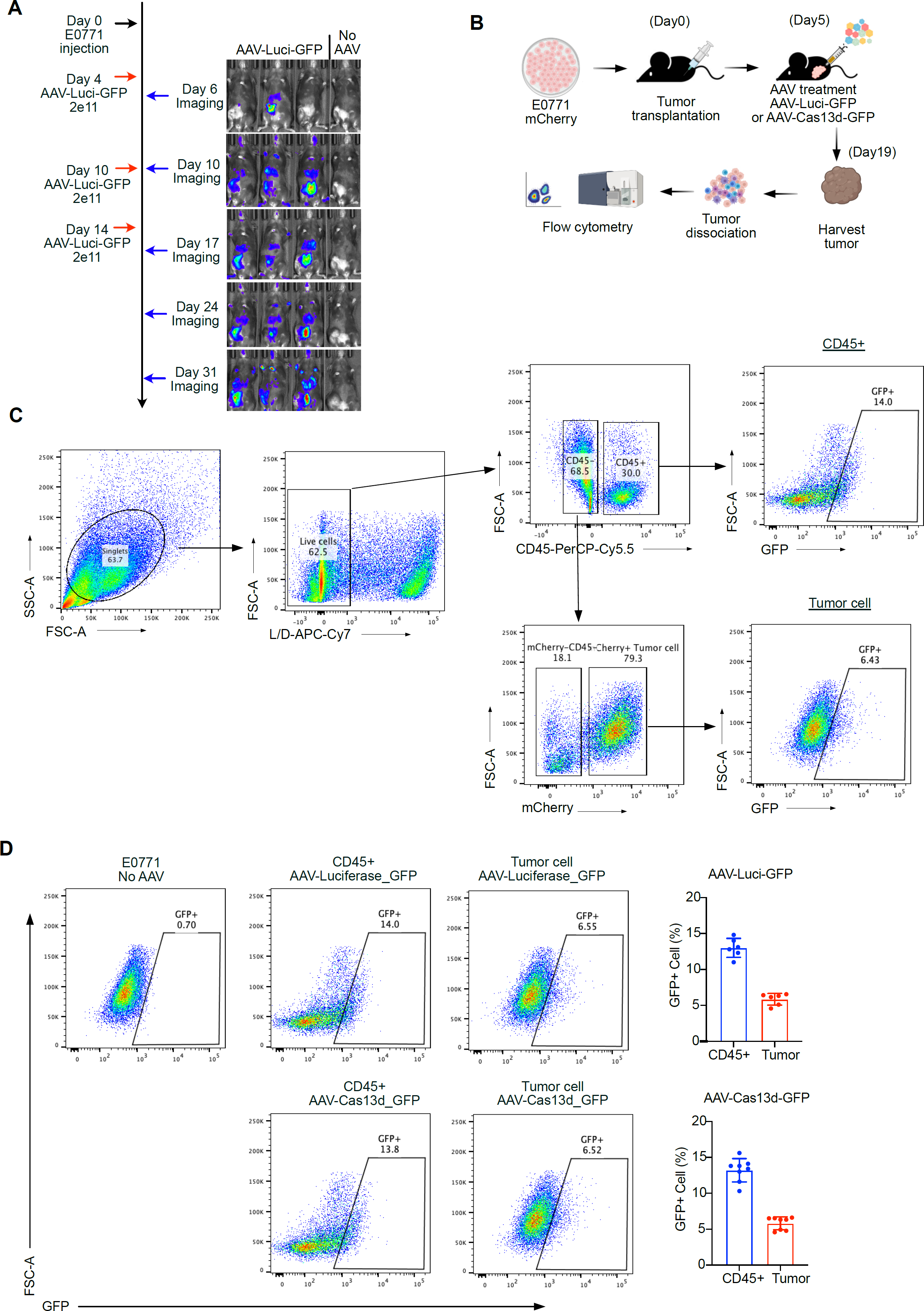
Persistent ectopic gene expression in tumors after intratumoral AAV injection. **A.** C57BL/6 mice were orthotopically injected with 2×10e^6^ E0771 cells. AAV-luciferase-GFP was then intratumorally injected at the indicated time points. In vivo bioluminescence imaging was performed to visualize luciferase activity. **B.** Schematic of experimental design for analyzing of AAV mediated gene expression in the tumor microenvironment (TME). AAV-Luciferase-GFP and AAV-Cas13d-GFP were used, and GFP expression in TME cell populations were analyzed. **C.** The gating strategy for flow cytometry staining panels are shown. Arrows indicate the parent population that the subsequent plot is gated on. **D.** GFP+ cells were gated as the AAV successfully infected cells. GFP+ was gated based on the E0771-mCherry cells without AAV infection.

**Figure S4.**
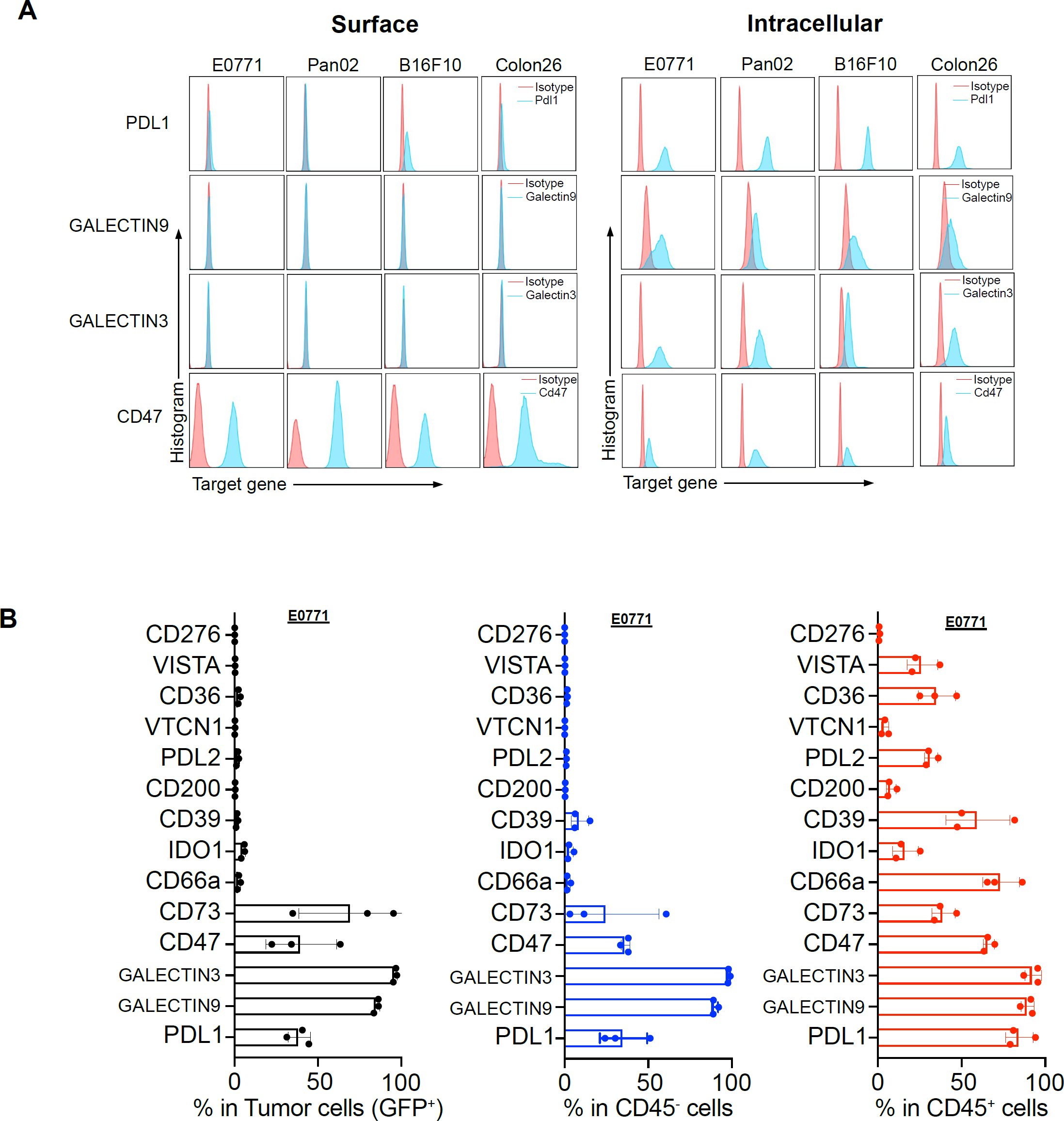
Flow cytometry analysis of immunosuppressive gene expression in cancer cell lines or in TME. **A.** Flow cytometry analysis of PGGC pool targets (PDL1, GALECTIN9, GALECTIN3, and CD47) in different murine cancer cell lines, either by surface or intracellular staining. **B.** Flow cytometry analysis of PGGC pool targets *in vivo* from syngeneic E0771 tumors. C57BL/6 mice (n = 3) were orthotopically injected with 2×e^6^ E0771-GFP cells. The tumors were harvested at 23 days post injection. Tumor tissues were dissociated for flow cytometry analysis of the indicated markers in each compartment.

**Figure S5.**
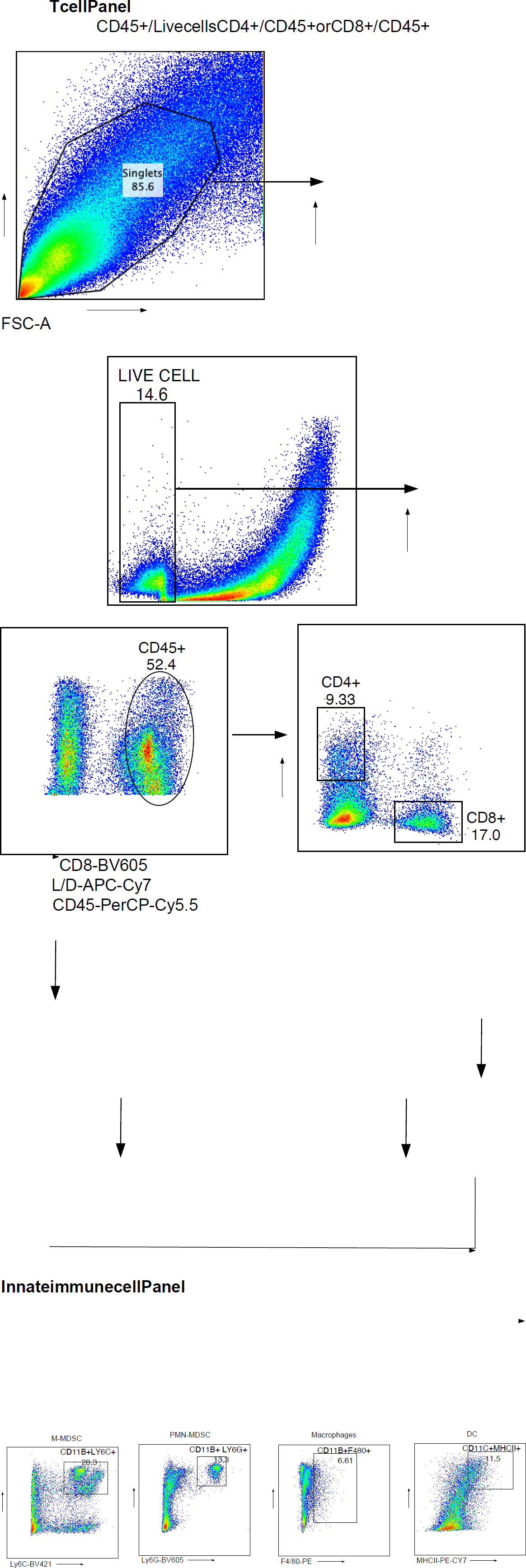
Gating strategy for the tumor infiltrating immune population analysis. The gating strategy for myeloid and lymphocyte cells flow cytometry staining panels are shown. Arrows indicate the parent population that the subsequent plot is gated on. CD8^+^ T = CD8^+^CD45^+^, CD4^+^ = CD4^+^CD45^+^, Macrophage = CD11b^+^F4/80^+^, Dendric cell (DC) = CD11c^+^MHCII^+^, MDSC = CD11b^+^Ly6G^+^(PMN-MDSC) + CD11b^+^Ly6C^+^(M-MDSC).

**Figure S6.**
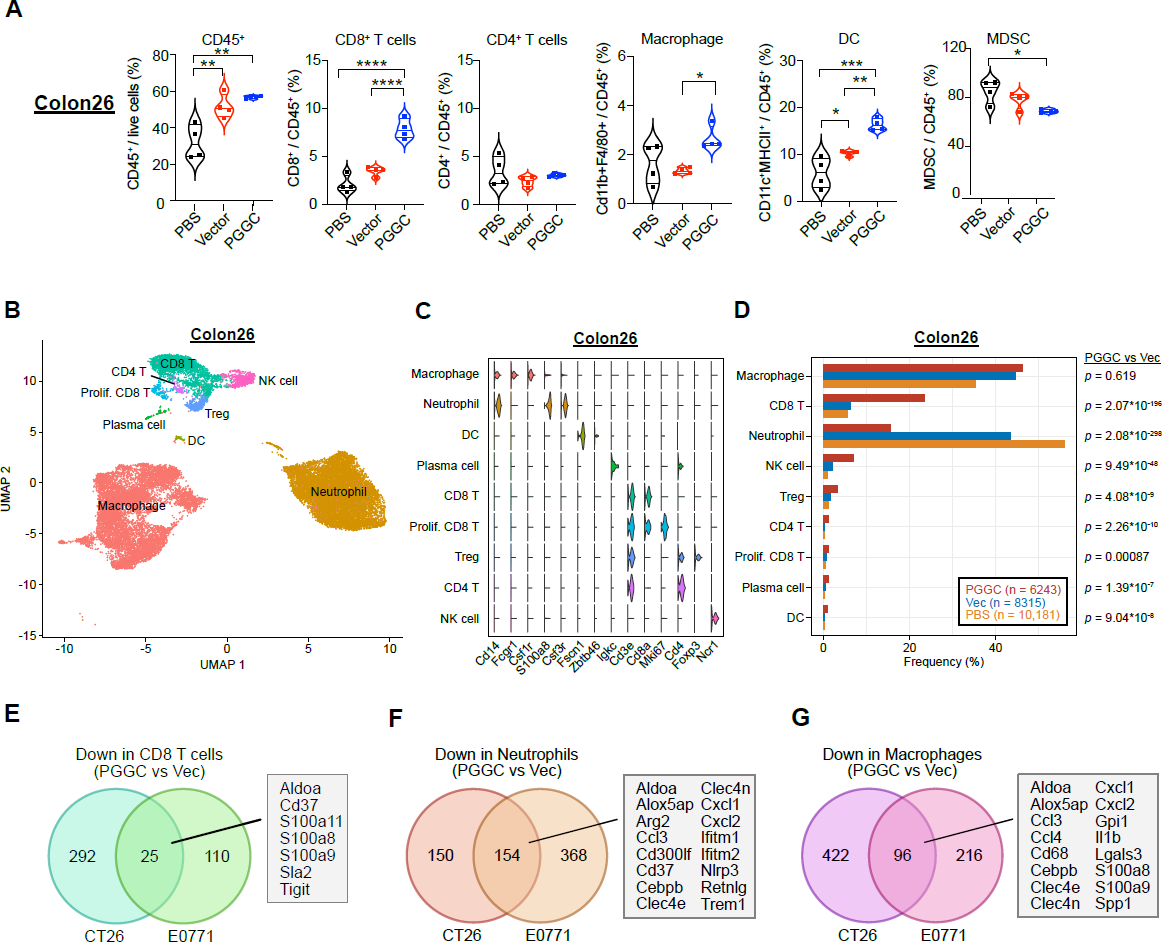
AAV-PGGC treatment remodels the immunosuppressive tumor microenvironment in CT26 colon cancer model. **A.** Relative abundances of several immune populations in subcutaneous Colon26 tumors. Mice were intratumorally treated with PBS (n = 4), AAV-Vector (n = 4) or AAV-PGGC (n = 4) at days 4, 9 and 14. Statistical significance was assessed by one-way ANOVA Tukey’s multiple comparisons test, adjusted P Value. (* *p* < 0.05, ** *p* < 0.01, *** *p* < 0.001). Non-significant comparisons not shown. **B.** UMAP visualization of single tumor-infiltrating immune cells, profiled by scRNA-seq. Mice bearing subcutaneous Colon26 tumors were treated with PBS, AAV-Vector or AAV-PGGC at days 4, 9 and 14. Tumors were harvested at day 29, and live CD45^+^ cells were sorted for scRNA-seq. **C.** Violin plots showing the expression levels of representative marker genes across the main cell clusters. **D.** Relative proportions of each cell type, across treatment groups. Statistical analysis between groups was performed by two-tailed Fisher’s exact test. **E-G.** Common signatures of downregulated genes in immune cell populations upon AAV-PGGC treatment from single cell data. Overlap of the down-regulated genes in CD8^+^ T cells (**E**), neutrophils (**F**) and macrophages (**G**), comparing AAV-PGGC vs AAV-Vector (Cas13d) in both the Colon26 and E0771 tumor models.

## Other Supplementary Materials for this manuscript include the following

### DNA oligonucleotide sequence information

All oligo sequences used in this study were listed in an excel file.

### Dataset S1 - Source data and statistics

Source data and statistics of non-NGS type data are provided in an excel file.

### Dataset S2 - NGS data

Processed data and statistics of NGS data are provided in an excel file.

## References

1. Rabinovich, G.A., Gabrilovich, D. & Sotomayor, E.M. Immunosuppressive strategies that are mediated by tumor cells. Annu Rev Immunol 25, 267–296 (2007).

2. Binnewies, M., et al. Understanding the tumor immune microenvironment (TIME) for effective therapy. Nat Med 24, 541–550 (2018).

3. Tormoen, G.W., Crittenden, M.R. & Gough, M.J. Role of the immunosuppressive microenvironment in immunotherapy. Adv Radiat Oncol 3, 520–526 (2018).

4. Kim, R., Emi, M., Tanabe, K. & Arihiro, K. Tumor-driven evolution of immunosuppressive networks during malignant progression. Cancer Res 66, 5527–5536 (2006).

5. Munn, D.H. & Bronte, V. Immune suppressive mechanisms in the tumor microenvironment. Curr Opin Immunol 39, 1–6 (2016).

6. Buchbinder, E.I. & Desai, A. CTLA-4 and PD-1 Pathways: Similarities, Differences, and Implications of Their Inhibition. Am J Clin Oncol 39, 98–106 (2016).

7. Pardoll, D.M. The blockade of immune checkpoints in cancer immunotherapy. Nat Rev Cancer 12, 252–264 (2012).

8. Wei, S.C., Duffy, C.R. & Allison, J.P. Fundamental Mechanisms of Immune Checkpoint Blockade Therapy. Cancer Discov 8, 1069–1086 (2018).

9. Sharma, P., et al. The Next Decade of Immune Checkpoint Therapy. Cancer Discov 11, 838–857 (2021).

10. Wolchok, J.D., et al. Nivolumab plus ipilimumab in advanced melanoma. N Engl J Med 369, 122–133 (2013).

11. Hammers, H., et al. Phase I Study of Nivolumab in Combination with Ipilimumab in Metastatic Renal Cell Carcinoma (Mrcc). Annals of Oncology 25(2014).

12. Rotte, A. Combination of CTLA-4 and PD-1 blockers for treatment of cancer. J Exp Clin Cancer Res 38, 255 (2019).

13. Sharma, P., Hu-Lieskovan, S., Wargo, J.A. & Ribas, A. Primary, Adaptive, and Acquired Resistance to Cancer Immunotherapy. Cell 168, 707–723 (2017).

14. Boettcher, M. & McManus, M.T. Choosing the Right Tool for the Job: RNAi, TALEN, or CRISPR. Mol Cell 58, 575–585 (2015).

15. Granados-Riveron, J.T. & Aquino-Jarquin, G. CRISPR-Cas13 Precision Transcriptome Engineering in Cancer. Cancer Res 78, 4107–4113 (2018).

16. Konermann, S., et al. Transcriptome Engineering with RNA-Targeting Type VI-D CRISPR Effectors. Cell 173, 665–676 e614 (2018).

17. Yan, W.X., et al. Cas13d Is a Compact RNA-Targeting Type VI CRISPR Effector Positively Modulated by a WYL-Domain-Containing Accessory Protein. Mol Cell 70, 327–339 e325 (2018).

18. Tang, X.E., Tan, S.X., Hoon, S. & Yeo, G.W. Pre-existing adaptive immunity to the RNA-editing enzyme Cas13d in humans. Nat Med 28, 1372–1376 (2022).

19. Wessels, H.H., et al. Massively parallel Cas13 screens reveal principles for guide RNA design. Nat Biotechnol 38, 722–727 (2020).

20. Shi, P., et al. RNA-guided cell targeting with CRISPR/RfxCas13d collateral activity in human cells. bioRxiv, 2021.2011.2030.470032 (2021).

21. Kelley, C.P., Haerle, M.C. & Wang, E.T. Negative autoregulation mitigates collateral RNase activity of repeat-targeting CRISPR-Cas13d in mammalian cells. bioRxiv, 2021.2012.2020.473384 (2021).

22. Wei, J., et al. Deep learning and CRISPR-Cas13d ortholog discovery for optimized RNA targeting. bioRxiv, 2021.2009.2014.460134 (2022).

23. Wang, D., Tai, P.W.L. & Gao, G. Adeno-associated virus vector as a platform for gene therapy delivery. Nat Rev Drug Discov 18, 358–378 (2019).

24. Ru, B., et al. TISIDB: an integrated repository portal for tumor-immune system interactions. Bioinformatics 35, 4200–4202 (2019).

25. Manguso, R.T., et al. In vivo CRISPR screening identifies Ptpn2 as a cancer immunotherapy target. Nature 547, 413–418 (2017).

26. Wang, X., et al. In vivo CRISPR screens identify the E3 ligase Cop1 as a modulator of macrophage infiltration and cancer immunotherapy target. Cell 184, 5357–5374 e5322 (2021).

27. Ishizuka, J.J., et al. Loss of ADAR1 in tumours overcomes resistance to immune checkpoint blockade. Nature 565, 43–48 (2019).

28. Shen, L., Evel-Kabler, K., Strube, R. & Chen, S.Y. Silencing of SOCS1 enhances antigen presentation by dendritic cells and antigen-specific anti-tumor immunity. Nat Biotechnol 22, 1546–1553 (2004).

29. Lawson, K.A., et al. Functional genomic landscape of cancer-intrinsic evasion of killing by T cells. Nature 586, 120–126 (2020).

30. Tang, Z., Kang, B., Li, C., Chen, T. & Zhang, Z. GEPIA2: an enhanced web server for large-scale expression profiling and interactive analysis. Nucleic Acids Res 47, W556–W560 (2019).

31. van Spriel, A.B., et al. A regulatory role for CD37 in T cell proliferation. J Immunol 172, 2953–2961 (2004).

32. Kong, Y., et al. T-Cell Immunoglobulin and ITIM Domain (TIGIT) Associates with CD8+ T-Cell Exhaustion and Poor Clinical Outcome in AML Patients. Clin Cancer Res 22, 3057–3066 (2016).

33. Grzywa, T.M., et al. Myeloid Cell-Derived Arginase in Cancer Immune Response. Front Immunol 11, 938 (2020).

34. Gómez-Herranz, M., et al. The effects of IFITM1 and IFITM3 gene deletion on IFNγ stimulated protein synthesis. Cell Signal 60, 39–56 (2019).

35. Srikrishna, G. S100A8 and S100A9: new insights into their roles in malignancy. J Innate Immun 4, 31–40 (2012).

36. Baghban, R., et al. Tumor microenvironment complexity and therapeutic implications at a glance. Cell Commun Signal 18, 59 (2020).

37. Ribeiro Franco, P.I., Rodrigues, A.P., de Menezes, L.B. & Pacheco Miguel, M. Tumor microenvironment components: Allies of cancer progression. Pathol Res Pract 216, 152729 (2020).

38. Motz, G.T. & Coukos, G. Deciphering and reversing tumor immune suppression. Immunity 39, 61–73 (2013).

39. Ni, G., et al. Targeting interleukin-10 signalling for cancer immunotherapy, a promising and complicated task. Hum Vaccin Immunother 16, 2328–2332 (2020).

40. Biswas, S., et al. Inhibition of TGF-beta with neutralizing antibodies prevents radiation-induced acceleration of metastatic cancer progression. J Clin Invest 117, 1305–1313 (2007).

41. Deyle, D.R. & Russell, D.W. Adeno-associated virus vector integration. Curr Opin Mol Ther 11, 442–447 (2009).

42. Joung, J., et al. Genome-scale CRISPR-Cas9 knockout and transcriptional activation screening. Nat Protoc 12, 828–863 (2017).

